# Jammed microgel growth medium prepared by flash-solidification of agarose for 3D cell culture and 3D bioprinting

**DOI:** 10.1101/2023.01.19.524686

**Authors:** M Sreepadmanabh, Meenakshi Ganesh, Ramray Bhat, Tapomoy Bhattacharjee

## Abstract

Although cells cultured in 3D platforms are proven to be beneficial for studying cellular behavior in settings similar to their physiological state, due to the ease, convenience, and accessibility, traditional 2D culturing approaches are widely adopted. Jammed microgels are a promising class of biomaterials well-suited for 3D cell culture, tissue bioengineering, and 3D bioprinting. However, existing protocols for fabricating such microgels either involve complex synthesis steps, long preparation times, or polyelectrolyte hydrogel formulations that sequester ionic elements from the cell growth media. Hence, there is an unmet need for a universally biocompatible, high-throughput, and widely accessible manufacturing process. We address these demands by introducing a rapid, high-throughput, and remarkably straightforward method to synthesize jammed microgels composed of flash-solidified agarose granules directly prepared in a culture medium of choice. Our jammed growth media are optically transparent, porous, yield stress materials with tunable stiffness and self-healing properties, which makes them ideal for 3D cell culture as well as 3D bioprinting. The charge-neutral and inert nature of agarose make them suitable for culturing various cell types and species, the specific growth media for which do not alter the chemistry of the manufacturing process. Unlike several existing 3D platforms, these microgels are readily compatible with standard techniques such as absorbance-based growth assays, antibiotic selection, RNA extraction, and live cell encapsulation. In effect, we present a versatile, highly accessible, inexpensive, and easily adoptable universal biomaterial for 3D cell culture and 3D bioprinting. We envision their widespread application not just in routine laboratory settings but also in designing multicellular tissue mimics and dynamic co-culture models of physiological niches.

## 1. Introduction

Cells grown as monolayers on conventional two-dimensional (2D) culture plates are significantly different from their *in vivo* counterparts in many different ways [1]. In standard flat plate cultures, not only do they exhibit altered morphologies and dynamics [2-4], their gene expression and signal transduction profiles are dramatically altered [5]. Furthermore, cells *in vivo* are exposed to more complex mechanical cues and chemical fields due to their three-dimensional (3D) packing and access to systemic circulation [6-15]. These mechanical cues from the microenvironment exert dramatic effects on processes as diverse as transcription, adhesion, migration, and metastasis and thus directly control cell fate specification [16-24]. Thus, a 3D culture system that replicates physiologically relevant conditions is essential for probing cellular functioning and behaviour *in vitro* [25]. Mimicking such scenarios requires biomaterials with tunable mechanical, chemical, and surface properties. Although, recent technological advances have addressed many of these requirements [26-30], achieving ubiquity and convenience using currently available 3D cell culture models still remain challenging.

Enabling the widespread adoption and effective utilization of 3D *in vitro* growth media imposes several demands on the design of such platforms. First, to support cell culturing across different time scales, a 3D medium with time-independent material properties is required. These 3D media should allow cell seeding, spatiotemporal probing, and endpoint retrieval of cells without destroying the structural integrity or altering the bulk properties of the matrix. Further, to match the mechanically diverse microenvironments encountered *in vivo*, 3D growth media with tunable stiffness is desirable. Additionally, visualization of cells deep inside the 3D growth media requires optically transparent substrata which can remain stable under standard culture conditions (typically, 30-37°C). Finally, such biomaterials should be broadly compatible across different cell types to support multi-tissue/multi-organ co-culture systems.

Jammed polyelectrolyte microgel systems—with their excellent material properties such as precisely tunable stiffness, optically translucent nature, and unimpeded nutrient transport—fulfil a significant fraction of these demands and have been extensively implemented for 3D cell culture and tissue engineering [31-36]. Additionally, such jammed microgel systems have self-healing material properties; they can transition from solid-like behavior to fluid-like behavior under applied shear [37]. This physical property has been effectively leveraged for freeform 3D bioprinting applications, allowing an injection nozzle moving through a jammed microgel system to locally shear the packing and inoculate cells in the inter-microgel space that get trapped in space once the nozzle is removed [37-43]. While these innovations have advanced our capabilities for 3D culture and tissue engineering, a handful of limitations still remain to be addressed. Commercially available microgels provide negligible control over the composition and charge density of the microgels whereas in-house fabrication of these microgels requires long preparation times with complex synthesis steps. Such caveats complicate their percolation into mainstream biological research.

Here, we present a versatile, cost-effective, highly accessible, and readily adaptable method for the rapid synthesis of jammed agarose microgels as a universal 3D biomaterial. The broad biocompatibility of agarose due to its inert and neutral nature makes it an ideal scaffold material. Micron-scale droplets of hot agarose suspension are flash-solidified directly in cold cell growth media to form agarose granules. Our one-step approach obviates several complexities of current microgel fabrication methods – such as dedicated microfluidic setups, complex organic synthesis, extensive wash steps, and trace levels of contaminants. A jammed packing of these agarose granules acts as a universal biomaterial that can support bacteria, yeast as well as mammalian cells in 3D. These jammed microgel media are translucent and enable embedded 3D bioprinting inside them due to their yield stress behavior and self-healing property. We demonstrate the suitability of these jammed agarose microgels for 3D *in vitro culture*, live cell imaging, antibiotic selection, 3D bioprinting, predetermined structure assembly and retrieval, as well as collagen-based functionalization. This dynamic range of biological capabilities promises a vast potential for tissue bioengineering and sculpting physiologically accurate replicas of diverse cellular microenvironments. Consequently, we believe this to be a valuable and readily adaptable tool for advancing ongoing research in domains such as mechanotransduction, durotaxis, and host-pathogen interactions, among others.

## 2. Materials and methods

### 2.1 Microgel Synthesis

Jammed agarose microgels are synthesized using agarose (HiMedia, MB002) as the base material. For a limited set of experiments (SI Fig. 2), agar from two different sources (Sigma, A1296 and Qualigens, Q21185) is also used to demonstrate the general applicability of our manufacturing method. Briefly, agar/agarose solutions of specified concentrations (w/v %) are prepared in double distilled water by suspending the powder form of agarose/agar in room temperature water, followed by heating the mixture in an autoclave to dissolve the solids. Briefly, one part of the warm (∼65°C) molten agar/agarose solution is rapidly injected at a rate of 1 ml/sec using a 0.5 inch-long, 26 gauge needle (Dispo Van) into four parts of continuous phase solvent (water, DMEM, RPMI, YPD, or LB) maintained within an ice-bath (0°C) with vigorous stirring at 1200 RPM. Injection at such a high flow rate, combined with the mechanical agitation, breaks up the molten agarose into micron-sized droplets, which are instantaneously solidified due to the rapid gelation of agar/agarose at low temperatures within the continuous phase. The resultant mixture is subsequently collected and centrifuged at approximately 7000Xg to yield two distinct phases - a densely packed microgel fraction that settles at the bottom and an overlying liquid solvent layer. The latter is gently decanted. Microgel fractions obtained in this manner are homogenized by gentle pipette mixing to ensure a smooth, uniform consistency.

Carbomer-based swollen granular hydrogels are prepared by dispersing 0.5% (w/v) Carbopol 980 polymer (Lubrizol, CBP1054A) in LB broth, stirring till complete dissolution, and adjusting the pH to 7 with 10N NaOH.

### 2.2 Characterization of jammed microgel media

#### 2.2.1 Rheological Properties

Rheological properties of the microgels are characterized using an Anton Parr MCR 302 rheometer. For all measurements, we use a roughened cone plate (CP50-4/S, diameter of 50 mm, cone angle of 3.987°, and cone truncation of 498 μm). In all cases, the sample is loaded onto the plate, following which the cone plate tool is gradually lowered until the sample becomes uniformly sandwiched between both surfaces, with the excess being trimmed away prior to measurement. Unless otherwise mentioned, all readings are obtained at room temperature. All our rheological measurements are shear-controlled, wherein we apply a known torque to the sample and record the consequent displacement of the measuring probe.

To evaluate the viscoelastic nature of jammed agarose microgels, we apply a low (1%) amplitude oscillatory shear strain over a range of frequencies while recording the the G’ (storage (elastic) modulus) and G’’ (loss (viscous) modulus). Consequently, for a viscoelastic material, G’ indicates the elastic solid-like behavior, while G’’ quantifies the viscous fluid-like nature. A G’ value greater than the G’’ indicates that the material predominantly behaves as a solid, while the alternate case implies a viscous fluid. Obtaining measurements across different frequencies helps interrogate the time-dependency (if any) of the material properties. Hence, the oscillatory frequency sweep tests help us establish the soft solid-like nature of our jammed agarose microgels under low shear regimes, which is also time-independent.

Separately, to understand the temperature-dependent gelation dynamics of agarose, we also apply the aforementioned rheological test under two different conditions. In one case, we maintain a sample of molten agarose (1% w/v) over a temperature gradient ranging from 65°C to 0°C by varying the temperature of the rheometer stage. Here, we record both the G’ and G’’ for oscillatory shear strain over different frequencies. Since we start off with a viscous solution, we expect the G’’ > G’. We track these two parameters while ramping down the temperature to 0°C and check for the transition point where the G’ first exceeds the G’’. This crossover point represents the temperature at which the molten agarose begins to gel, and thereby transitions to a soft solid. The second case is where we maintain the rheometer’s measuring stage at 0°C, load a warm molten solution of 1% w/v agarose, and immediately commence measuring the G’ and G’’ as described above. This test helps characterize the kinetics of the gelation process. We observe that the experimental setup time, which is of the order of a few seconds, is sufficient for the agarose to gel. Hence, we do not observe a crossover point where G’ > G’’. Rather, the G’ remains higher than the G’’ from the first timepoint itself, indicating the rapid, almost instantaneous gelation of agarose upon contact with a cold surface.

Several previously reported jammed microgel systems have been shown to exhibit yield stress behavior. Briefly, these microgels are fluidized upon application of high shear rates while reversibly transitioning to a soft solid-like nature under lower shear. This is an important property for 3D printing. The fluidizable nature is necessary for the printing nozzle to freely translate within the microgel bath, while the solid-like properties are required to stably hold the extruded structures in place. To check for the yield stress nature of our jammed agarose microgels, we apply a unidirectional shear at varying rates (30 seconds for each) while recording the shear stress response. At higher shearing rates, we observe a linear relation between the applied shear and the recorded stress. However, a plateauing trend for the stress response below a certain shear rate indicates the presence of yield stress behavior. This is termed the crossover shear rate, i.e., the shear below which the material is a soft solid and above which it is fluidized. These observations confirm the yield stress material properties of agarose microgels.

#### 2.2.2. Particle size distributions

To determine microgel particle sizes, a 1:1000 dilution of the microgels is prepared in water. Particles are imaged using a phase contrast microscope (Nikon Eclipse TE300) under 10X magnification. These images detail 2D projections of three-dimensional microparticles. The micrographs are analyzed in ImageJ by manually segmenting the particle outlines and employing built-in functions for measuring the particle area and perimeter. For our present study, two parameters have been quantified. The first is the square root of the area, which is an approximate indicator of the particle size. To understand the variability in particle shape, we also calculate the shape factor—the perimeter divided by the square root of the area.

#### 2.2.2. Pore size distributions

To determine the inter-particle pore spaces, we use 200 nm fluorescent tracer particles and track their thermal diffusion through the inter-particle pore spaces. Since the agarose microparticles are rigid bodies, the tracer particles only traverse the pores. We disperse a dilute solution of these in jammed 1% agarose microgels prepared in LB and track their thermal diffusion-guided trajectory by acquiring images within a time interval of 17 ms using the 40X objective of an inverted laser-scanning confocal microscope (Nikon A1R HD25). Mapping the center of each particle over time using a peak-finding function, we obtain their MSD (mean square displacement). The underlying assumption is that over short time scales, the particles may freely explore the pore spaces, with their motion being constricted or altered only via contact with the surfaces circumscribing each pore. Consequently, the particle MSDs should increase linearly with time for such unimpeded motions while plateauing upon encountering the pore’s boundaries. The square root of these plateau values added to the particle diameter gives the characteristic smallest pore space dimensions explored by the particles, the 1-CDF (cumulative distribution function) of which we report as the overall pore size distribution for the LB-based 1% agarose microgel.

### 2.3. In vitro Experiments

Microgels synthesized using a 1% w/v agarose solution are used for all *in vitro* experiments. Since each cell type has a specific culture media composition optimal for its growth, we vary the continuous phase solvent in the manufacturing process as per requirement. Briefly, bacterial cells are cultured in 2% (w/v) LB (Luria-Bertani) broth (Sigma L3022); yeast cells are grown in YPD (Yeast-extract Peptone Dextrose, with 1% w/w yeast extract (Himedia RM027), 2% w/w tryptone (Himedia RM014), and 2% w/w D-glucose (Qualigens G15405)); and mammalian cells are grown in either DMEM (Dulbecco’s Modified Eagle Medium (Gibco, 12100-038), with 10% fetal bovine serum (HiMedia RM10432) and 1% Penicillin-Streptomycin) or RPMI (Roswell Park Memorial Institute Medium (HiMedia AT060), with 20% fetal bovine serum and 1% Penicillin-Streptomycin) culture media.

#### 2.3.1 Cell Culture

OVCAR-3, MDA-MB-231, hTERT FT 282, MeT-5A, and NIH-3T3, obtained from ATCC and used for mammalian cell culture experiments are maintained as adherent monolayers in tissue-culture-treated plastic plates. We also employ two different variants of the OVCAR-3 cells – one which stably expresses the green fluorescent protein (GFP) and another which expresses the red fluorescent protein (RFP) [44]. These are grown using DMEM (for MDMB-231, FT-282, MET5, and NIH-3T3) or RPMI (for OVCAR-3) complete culture media at 37°C under 5% CO2. In all cases, cells are harvested when cultures are 70-80% confluent, and manually seeded into the agarose microgels prepared using DMEM/RPMI complete culture media as the continuous phase. These samples are housed in 35-mm glass-bottom tissue culture dishes. The dishes have a 15-mm central circular cavity, to the base of which a cover glass slide is attached to generate glass bottom dishes for optimal imaging quality.

To culture yeast in jammed microgel media, we use *Saccharomyces cerevisiae*, a widely-studied species of yeast that reproduces by budding. We first prepare an overnight liquid culture in 2 ml of YPD using a 20 μl inoculum from frozen glycerol stocks (1:100 dilution). The culture is grown for 20 hours at 30°C while shaking the sample at 220 RPM until it attains the stationary phase. For subsequent live-cell imaging and growth curve experiments, a small volume (20 ul) from the overnight-grown culture is homogeneously mixed with 980 μl of YPD-based jammed agarose microgels (1:50 final dilution).

Additionally, the present work employs three strains of *Escherichia coli*. These are – a motile wild-type strain, a non-motile strain, and a strain carrying a kanamycin resistance-conferring plasmid. All three bacterial strains constitutively express the green fluorescent protein (GFP) throughout their cytoplasm. To culture bacteria in jammed agarose microgels, we first grow an overnight liquid culture in 2 ml of LB broth using a 20 μl inoculum from frozen glycerol stocks (1:100 dilution). These cultures are grown for 20 hours at 30°C while shaking at 220 RPM until they attain the stationary phase. For live-cell imaging, tracking, and growth curve assays, 10 μl of the overnight-grown culture is added to 990 μl of LB-based jammed agarose microgels (1:100 final dilution).

#### 2.3.2 Collagen-infused microgels

To manufacture adherent microgels, we fabricate jammed agarose microgels infused with polymerized Type 1 collagen fibrils within the inter-particle pore spaces. For this, we add a solution of acidic rat tail Type 1 collagen monomers (Invitrogen, A1048301) to pre-made jammed microgel growth media and initiate the polymerization process. Since collagen monomers are suspended within the microgel pores, the fibers formed post-polymerization will also be present in these regions. We homogeneously mix 200 μl of 1% RPMI-based agarose microgel, 2.5 μl of 1N NaOH, and 100 μl of 3 mg/ml Collagen Type I. Immediately following this, cells are dispersed within the sample by pipette mixing and incubated overnight at 37°C, 5% CO2. The polymerized collagen is visible as fibrous structures within the pore spaces, which we observe using reflectance with confocal microscopy.

#### 2.3.3 Cell Viability

To evaluate the viability of various cell types in DMEM/RPMI-based agarose gels, we employ calcein-AM and propidium iodide (PI)-based staining. Briefly, cleavage of the acetoxymethyl by intracellular esterases releases a fluorescent calcein molecule, which marks live cells. PI labels dead cells with compromised membranes by intercalating with the DNA and fluorescing. We stain adherent culture plates with calcein-AM (ThermoFisher Scientific, C1430) at a 1 μg/ml final concentration by resuspending from a 1 mg/ml stock in the appropriate culture media. Plates with the staining solution are incubated at 37°C, 5% CO_2,_ for 30 minutes, followed by replacement with fresh media and a recovery period of 1 hour at 37°C, 5% CO_2_. Following this, we harvest the cells and resuspend them in DMEM/RPMI-based agarose microgels. This mixture is subdivided into two fractions for time point-specific imagings at t = 0 hours and t = 24 hours. Prior to imaging, the cell-laden microgel samples are stained with PI (HiMedia, TC252) by directly adding the dye to achieve a final concentration of 5 μg/ml, followed by incubation for 15 minutes at 37°C, 5% CO2. While imaging, identical optical settings are maintained for both t = 0 hours and t = 24 hours samples, following which the total number of live and dead cells are manually tallied for each field imaged.

#### 2.3.4 Setup for Live Cell Imaging

We use an inverted laser-scanning confocal microscope (Nikon A1R HD25) for all image acquisitions. Specimens for imaging are housed in glass bottom 35 mm dishes. To minimize sample loss due to evaporation, these are covered with a thin film of mineral oil (Sigma, M5904), which allows oxygen to diffuse freely. For live-cell imaging, a stage-top incubator is used to maintain the optimal temperature (30°C for both yeast and bacterial cells, 37°C for mammalian cells).

For time-lapse imaging of yeast growth, we focus on individual yeast colonies using a 40 X objective and acquire images every 10 minutes. For time-lapse imaging of bacterial growth, we maintain a field of view with several visible bacteria in focus and acquire images every 60 minutes. Tracking of individual motile bacteria within the inter-particle pore spaces of the microgel is carried out using a 40X objective with 8X optical zoom, with a temporal resolution of 100 ms between successive frames. For mapping the trajectory of these bacteria, we pseudocolor individual bacterium within a field of view and manually track their center over successive time frames using an ImageJ-inbuilt function for particle tracking.

#### 2.3.5 Growth Curves

Well-grown stationary phase cultures of yeast and bacteria, obtained as described in section 2.3.1, are used as the inoculums for growth curve assay samples. For yeast, 20 μl of overnight liquid culture is added to 980 μl of sterile microgel; whereas for bacteria, 10 μl of overnight liquid culture is added to 990 μl of sterile microgel (1:50 and 1:100 dilution, respectively). Mixing is carried out carefully to ensure even dispersion while avoiding the formation of bubbles, as these interfere with the absorbance readings. 200 μl of well-mixed samples are added to separate wells of a 96-well plate, once again exercising caution to avoid forming bubbles. A simple strategy is to extrude a hanging droplet from the pipette tip, touch this to the bottom of the well, and gently dispense the rest of the sample. To prevent the gels from drying out, considering the minute volumes involved, double distilled water is added to the vacant wells surrounding the sample. Plates prepared in this manner are incubated at 30°C overnight in an automated multi-well plate reader, programmed to acquire absorbance readings for 600 nm at regular intervals (in our case, once every 10 minutes). These represent the optical density of the sample over time, which corresponds to the total growth and increase in biomass. Antibiotic treatments are carried out by adding kanamycin (HiMedia RM210) to the microgel samples to achieve a final concentration of 50 μg/ml.

#### 2.3.6 Encapsulation Protocol

To encapsulate bacteria within agarose microparticles, 2 ml of a well-grown stationary phase culture of GFP-expressing kanamycin-resistant *E. coli* is added to 50 ml of molten 1% w/v agarose solution prepared in LB broth. We prefer to use antibiotic-resistant E. coli as opposed to the wild-type strain for this particular experiment in order to minimize the potential for contamination during the gel synthesis and centrifugation steps. This is mixed well and rapidly to ensure even dispersion before the molten agarose commences gelation. The subsequent protocol is identical to what has been described above to generate jammed agarose microgels, with the continuous phase solvent being LB broth. The resultant microgel particles contain embedded bacteria, as is observed using microscopy.

#### 2.3.7 RNA Extraction, cDNA Synthesis, and qRT-PCR

For bacterial RNA extraction, the workflow described herein has been broadly based on a previously outlined method [45]. To prepare samples, we first grow overnight cultures of wild-type *E. coli*, as described above in section 2.3.1. A 20 μl inoculum from this is added to 5 ml of LB-based jammed agarose microgel. These samples are cultured at 30°C for 20 hours with shaking at 220 RPM until they attain the stationary phase. First, Triton X-100 is added to achieve a final concentration of 0.2%, with thorough mixing to ensure uniform dispersion. Samples are then subjected to heat treatment for two hours within a dry bath maintained at 65°C to ensure efficient cell lysis. This is rapidly followed by the addition of two parts chloroform and one part methanol per one part of the microgel specimen. Samples are mixed by inverting several times (it is advisable to avoid vortexing at this, as well as all subsequent stages, to avoid shearing the RNA), followed by centrifugation at 4°C and 7000 xg for 15 minutes. The resultant mixture shows a clear separation into three phases - an upper aqueous phase, an opaque interfacial layer, and a lower organic phase. The aqueous fraction, which contains the RNA, is carefully collected in a separate tube. To this, we add an equal volume of chloroform, mix by inverting, and centrifuge at 4 °C and 7000 xg for another 15 minutes. Once again, the aqueous phase obtained is transferred into a separate tube. In order to precipitate the RNA from this, 2 volumes of absolute ethanol and 1/10 volume of 3M sodium acetate (pH 5.2) are added per 1 volume of the collected aqueous phase. This is mixed well by gentle inverting of the tubes and incubated at -20°C for two hours to promote precipitation of the RNA fraction. Following this, the samples are centrifuged at 4 °C and 7000 xg for 15 minutes, followed by washing the resultant pellets with absolute ethanol under similar conditions. The final pellet obtained is then air-dried and resuspended in 100 μl of cold nuclease-free water, which is further centrifuged at 4 °C and 7000 xg for 15 minutes. Without disturbing the pellet (if any), the aqueous phase from this step is carefully collected in a separate tube. This represents the final extracted RNA fraction. The sample’s RNA concentration is determined using the Qubit Broad Range RNA Assay (Invitrogen, Q10210).

To create 3D microtissues, we first bioprint a homogeneous mixture of NIH 3T3 cells (20 μl cell pellet), collagen type I (10 μl of a 3 mg/ml stock), and 0.1 NaOH (2.5 μl) inside a jammed agarose microgel growth medium. We allow the tissue to stabilize for at least 24 hours under standard cell culture conditions (37°C, 5% CO_2_). To extract the RNA, the microtissue fragment is then picked up, washed with PBS, and digested with 1 ml of Trizol (RDP Trio Reagent, HiMedia, MB566). 200 μl of chloroform is added to the Trizol-digested sample, followed by vigorous mixing. Following centrifugation at 4°C and 12,000 xg for 10 minutes, we obtain three distinct phases – a clear aqueous phase on top containing the RNA, an interfacial layer in the middle with the DNA, and an organic phase comprising proteins at the bottom. The aqueous phase is carefully collected in a separate tube, followed by the addition of 500 μl isopropanol and mixing by gentle inversion and incubation at room temperature for 10 minutes. After centrifugation at 4°C and 12,000 xg for 10 minutes, we obtain a pale off-white pellet at the bottom. The supernatant is decanted, and the pellet is washed once with 75% ethanol in water by centrifuging at 4°C and 12,000 xg for 10 minutes. Once again, the supernatant is decanted, and the pellet is subjected to a dry spin by centrifuging under identical conditions as mentioned above. Finally, the pellet is air-dried to remove all residual traces of ethanol and resuspended in 30 μl of nuclease-free water. This solution contains the extracted total RNA, the concentration of which is determined using the Qubit Broad Range RNA Assay.

We prepare cDNA using samples of total RNA extracted from 3D microtissue as described above using the Superscript III First Strand (Invitrogen, 18080-051) kit. For this, 1 μl of a 50 μM oligo(dT)_20_ solution and 1 μl of a 10 mM dNTP mix are added to 8 μl of RNA (total amount of 5 μg). The mixture is incubated at 65°C for 5 minutes and then placed on ice for 1 minute. Subsequently, we prepare 10 μl of a cDNA synthesis mix by combining 2 μl of 10X RT buffer, 4 μl of 25 mM MgCl2, 2 μl of 0.1 M DTT, 1 μl of RNaseOUT (40 U/μl), and 1 μl of SuperScript III Reverse Transcriptase (200 U/μl). This is added to the mixture of total RNA and primers and incubated at 50°C for 50 minutes, followed by termination of the reaction by incubation at 85°C for 5 minutes and storage on ice. Finally, 1 μl of RNaseH is added to the mixture and incubated at 37°C for 20 min. The resultant solution is directly used as a template for subsequent PCR amplification.

Expression levels are evaluated using gene-specific primers for the PER2 (Period Circadian Regulator 2), BMAL1 (Brain and Muscle Arnt-like Protein-1), and DBP (D-Box Binding PAR BZIP Transcription Factor). From the cDNA prepared as described above, 3 μl of a diluted stock was added such as to achieve a final concentration of 15 ng per well in a 384-well plate. This serves as the template for the PCR amplification, which was carried out using the iTaq Universal SYBR Green Supermix (BioRad, 1725121), as per the manufacturer’s instructions. The reactions were carried out in an AppliedBiosystems ViiA 7 Real-time PCR System, using the in-built QuantStudio Real-time PCR software for defining the run protocol and data generation.

### 2.4 3D bioprinting in agarose microgels

During the initial course of this study, we noticed that 3D printing into jammed agarose microgels prepared as per the protocol described in section 2.1.1 resulted in coarse-grained structures without sharp, precisely-defined contours. While this is scarcely an impediment for elementary geometries, we identify this as a potential limitation towards achieving high-quality printing. We circumvent this by generating fine-grained microgels of smaller particle sizes for 3-D bioprinting by shearing microgels prepared as described above using a tissue homogenizer (Polytron PT2100). Due to their lower yield stress, these homogenized microgels allow for smoother translation of the 3D printing nozzle. This treatment generates a large number of air bubbles which are easily eliminated with a quick centrifugation step (short spin to 5000 xg), followed by pipette mixing to ensure a homogeneously smooth sample.

For bioprinting with bacteria, overnight cultures grown as described above are harvested by centrifuging at 5000 xg for 5 minutes to obtain a dense pellet, which is then loaded into a 1 mL syringe attached to a 20 gauge needle. Leveraging custom-designed structural supports, the syringe is loaded onto the pump arm of a 3-D printer (Newport, Motion Controller, Model XPS-D). All structures shown are printed using custom-written MATLAB codes to operate the printer. For bioprinting with OVCAR-3 cells, we harvest the cells by trypsinization, collect them as a dense pellet, and proceed in a manner identical to what has been described above for bacterial bioink, except that these are manually injected using a pipette into a bacteria-laden microgel. For retrieving 3-D printed structures from the bulk gel, 20 gauge needles attached to a 1 ml syringe are used to selectively extract the element of interest without perturbing the surrounding regions. This is achieved by holding the needle tip vertically above the specific region and drawing up the syringe plunger at a slow, steady pace. Post-printing, the microgel samples are layered on top with mineral oil to prevent desiccation.

## 3. Results and discussion

### 3.1 Synthesis of a neutral jammed microgel system

Several routine laboratory protocols in life science research, such as gel electrophoresis[46], microbial cultures[47], and surface coating[48, 49], employ a variety of chemically inert and porous hydrogel substrates made of crosslinked agarose. The gelation of agarose initiates when its monomeric units, D-galactose and 3,6-anhydrous-L-galactopyranose, crosslink through hydrogen bonding[50]. At high temperatures, disruption of these non-covalent H-bonding produces a viscous agarose solution. At low temperatures (below ∼35°C), an aqueous solution of agarose transiently gels to form a solid monolithic matrix. Interestingly, this liquid-to-gel transition of agarose shows thermal hysteresis: while the gelling initiates at ∼ 35°C temperature, the melting of agarose gels starts at a much higher temperature (∼85°C)[51]. Thus, after gelation, solid agarose gels remain thermally stable at standard culture temperatures. Furthermore, agarose gels have tunable material properties; the concentration of agarose in these hydrogels regulates their mechanical properties as well as the pore sizes between agarose fibers[52]. Typical pore sizes are much larger (∼100 nm) than nutrient molecules but smaller than most organisms, which allows for unimpeded nutrient transport while restricting cells to the surface[53]. Hence, these characteristics, in combination with their neutral and inert nature, render agarose gels highly compatible with several biomedical applications ranging from tissue regeneration for drug delivery and live-cell encapsulation [54].

Drawing our inspiration from the wide application of agarose in biological sciences, in the present study, we introduce a high-throughput flash solidification method to prepare micron-scale granules of agarose hydrogels by leveraging the temperature-dependent gelation process. To quantify the gelation dynamics, a warm solution of agarose (65°C, 1wt% solid) is loaded between two parallel plates of a rheometer, on which we perform two different sets of rheological tests. First, we apply a small amplitude (1%) of oscillatory strain at 1 Hz frequency and measure the storage (elastic) modulus (G’) and loss (viscous) modulus (G’’) as a function of plate temperature. We find that at high-temperatures, G’’ is much higher than G’, which is a key property of viscous fluids and indicates that the H-bonding within agarose is yet to initiate. Non-covalent H-bonding interactions increase with the continuous decrease in temperature, thereby increasing the solid-like behavior. Indeed, beyond a crossover temperature (∼30°C), G’ exceeds G’’ when the agarose solution gels into a solid matrix (SI Fig. 1A). Next, we enhance the rate of gelation by dramatically widening the temperature difference between the hot agarose solution and cold surroundings. For these tests, we directly inject the hot agarose solution between two parallel plates of a rheometer maintained at 0°C and immediately commence the measurements of G’ and G’’(SI Fig. 1B). This whole procedure requires less than 10 seconds: long enough for the instantaneous gelation of the agarose solutions upon contact with the cold surfaces. This rapid liquid-to-solid phase transition is the cornerstone of our microgel synthesis strategy.

More precisely, an autoclaved aqueous suspension of agarose above the gelation temperature (∼65°C) is injected via a 26 gauge syringe needle into an ice-cold water bath (0°C) at a high flow rate of ∼1 ml/s. Injection at such a high flow rate produces turbulent flow at Reynolds number ∼ 5000. Simultaneously, the ice-cold water bath is vortexed at 1200 RPM to maintain a turbulent regime. The vigorous and turbulent shear flow breaks up the cold and solidified agarose from the agarose solution and forms micron-scale granules (Fig. 1A). Maintaining a constant bath to final solid volume fraction, injection at a preferred Reynolds number, precise agarose concentration, controlled vortexing speed, and bath temperature, ensure precise particle sizes. To characterize this, we image individual microgel particles and measure their size. The average particle sizes are very consistent across different batches and are in the order of 100 +/-50 μm (Fig. 1B, SI Fig. 2A, and SI Fig. 2D). Subsequently, these microgels are randomly packed into a jammed state *via* centrifugation, and the supernatant fluid is removed to form a three-dimensional matrix.

**Figure 1.**
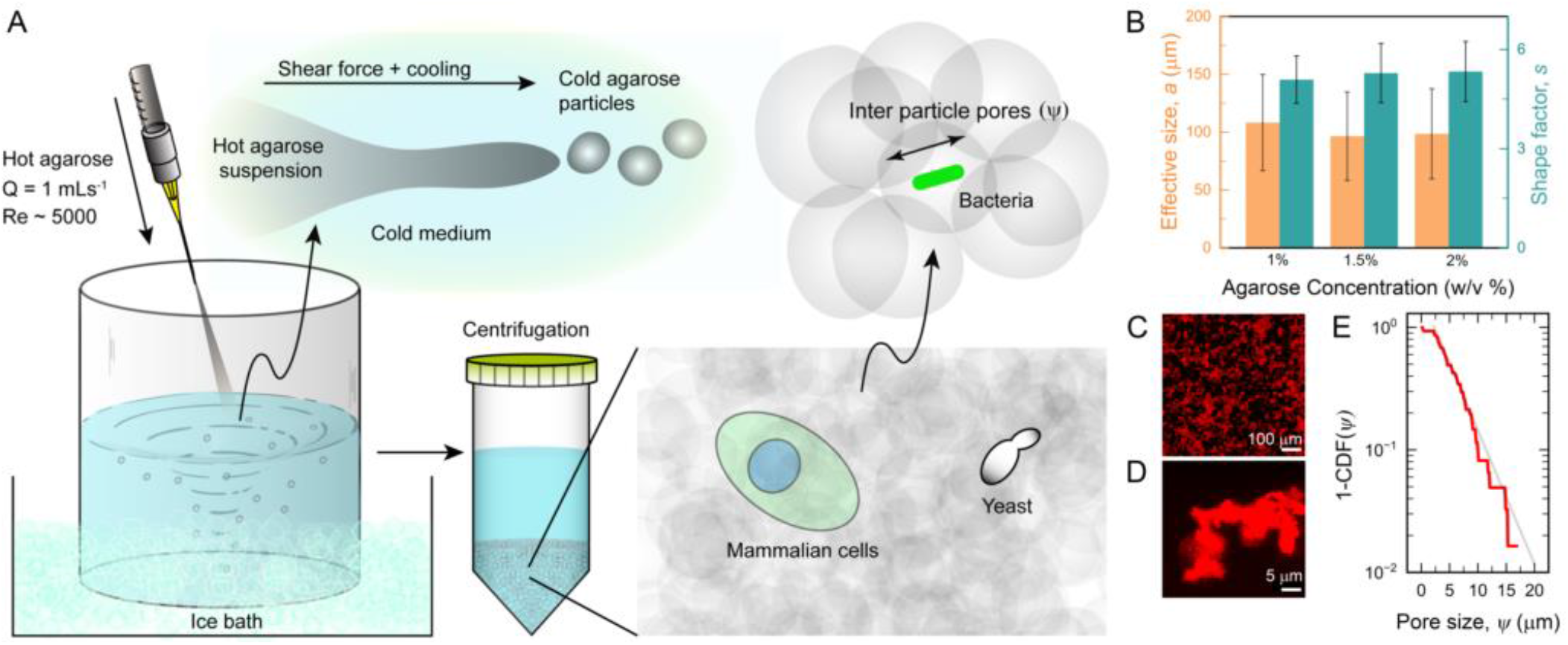
Jammed agarose microgels for 3D cell culture. (**A**) The synthesis of these microgels is carried out using a flash solidification technique which involves rapid gelation of micron-scale molten agarose droplets within a reservoir of media maintained at low temperatures and high Reynolds number. The granules thus generated are collected and assembled into a jammed packing by centrifugation to obtain a 3D growth media. Cellular proliferation within these jammed microgel-based growth media essentially occurs in a suspended 3D environment. While single microorganism cells reside within the inter-particle pore spaces, large mammalian cells as well as microorganism colonies rearrange the particles locally to migrate, proliferate, and expand. (**B**) The manufacturing process of microgels conserves the particle size and shapes. Regardless of the concentration of agarose used, the particle sizes and shape factor remain consistent. (**C**) We visualize the pore spaces by allowing fluorescent tracer particles to freely diffuse through the pore space and imaging the tracer locations over time. A time projection of all possible tracer locations for 10 minutes shows the disordered nature of the pore space. (**D**) Image of an individual pore confirms the large void volume of the 3D media. (**E**) By tracking the thermal diffusion of individual tracer particles, we measure the pore size—the smallest dimension of the pore—and plot the cumulative distribution the pore sizes.

To culture cells in this 3D medium, we need to ensure a smooth transportation of oxygen and nutrients molecules to the cells. In this case, the diffusive transport of nutrients and oxygen is governed by the porosity of the jammed microgel medium. Although the pores inside the individual microgel particles are large enough for nutrient diffusion, there exists another set of inter-microgel pores, which are orders of magnitude larger in size. To quantify the inter microgel pore sizes, we deploy tracer particles and image their motion under thermal diffusion. A time projection (∼10 min) of all possible positions of the tracer particles indicates the tortuous pore spaces inside the jammed microgel medium (Fig. 1C). To further characterize the smallest dimensions of these pores, we track the centroid of these tracer particles and measure the cage size from their mean squared displacement (see materials and methods). Similar to many other randomly packed porous media, we find these pore sizes are exponentially distributed (Fig. 1E). By fitting an exponential function to this data, we estimate an average pore size of 4 μm. Presence of such large pores—which are several orders of magnitude larger than most small molecules—ensures unimpeded supply of nutrients that is critical for the growth and metabolism of cells (Fig. 1E).

Microgel production *via* flash solidification of agarose solutions overcomes a majority of the complexities of state-of-the-art microgel manufacturing methods. Current practices of microgel synthesis employ either a bottom-up or top-down approach [55]. In the bottom-up approach, micron-sized crosslinked hydrogel particles are made from a precursor solution either through a precipitation polymerization method—wherein both the monomer and catalytic initiator are soluble, but the insoluble polymeric end product precipitates out—or emulsion polymerization—wherein emulsified droplets of aqueous monomers solutions are polymerized and crosslinked inside an oil bath. However, hydrogel particles produced in these methods require the removal of the solvents and oils through multiple washing and purification steps. In top-down approaches, hydrogels are mechanically sheared into smaller particles, but this requires equipment-specific starting volumes and multiple processing steps. Other drawbacks include long preparation times, low optical transparency, and the presence of tiny fragments of polymer chains that increase the osmotic pressure of the medium. Our approach neatly circumvents these problems by directly injecting hot agarose solutions into an ice-cold liquid culture medium of choice. This ensures we have no unwanted residuals since the only reagents involved are agarose and the growth media itself. Micro-particles thus formed are recovered by centrifugation, yielding a ready-to-deploy jammed microgel system without any downstream processing steps. The supernatant growth medium removed after centrifugation is easily reusable and hence reduces the wastage of cell growth media in the washing process.

### 3.2 Characterization of jammed agarose microgels

For long-term cell culturing practices, it is critical for the 3D growth matrix to keep the cells suspended in space— ideally for several days—and allow them to form multi-cellular populations. Although several suspension culture media have been prepared from viscous fluids and utilized to culture cells in 3D, they fail to provide long time stability to individual cells and multicellular structures; suspended cellular populations eventually settle down over time. Thus, a 3D growth medium is preferred to have solid-like behavior. To test if our jammed agarose medium has a solid-like material property, we perform rheological measurements where we apply a small amplitude of oscillatory shear strain (1%) at varying frequencies (1Hz to 0.1Hz) and measure the shear moduli of the packings. Multiple batches and combinations of jammed agarose media were tested. In all measurements, the storage (elastic) modulus (G’) remains frequency independent and higher than the loss (viscous) modulus (G’’) —a characteristic behavior of a solid-like material. These measurements confirm that within elastic limits, jammed agarose microgels behave like soft solids with time-independent material properties (Fig. 2A, SI Fig. 2B, and SI Fig. 2E). Furthermore, a prime motivation for developing 3D culturing systems is to accurately mimic physiological microenvironments with tunable mechanical properties. By regulating the agarose concentration, we control the overall stiffness of the jammed microgels. This tunable stiffness of our microgel system, along with its temporal stability, makes it a reliable scaffold for *in vitro* cell culture.

**Figure 2:**
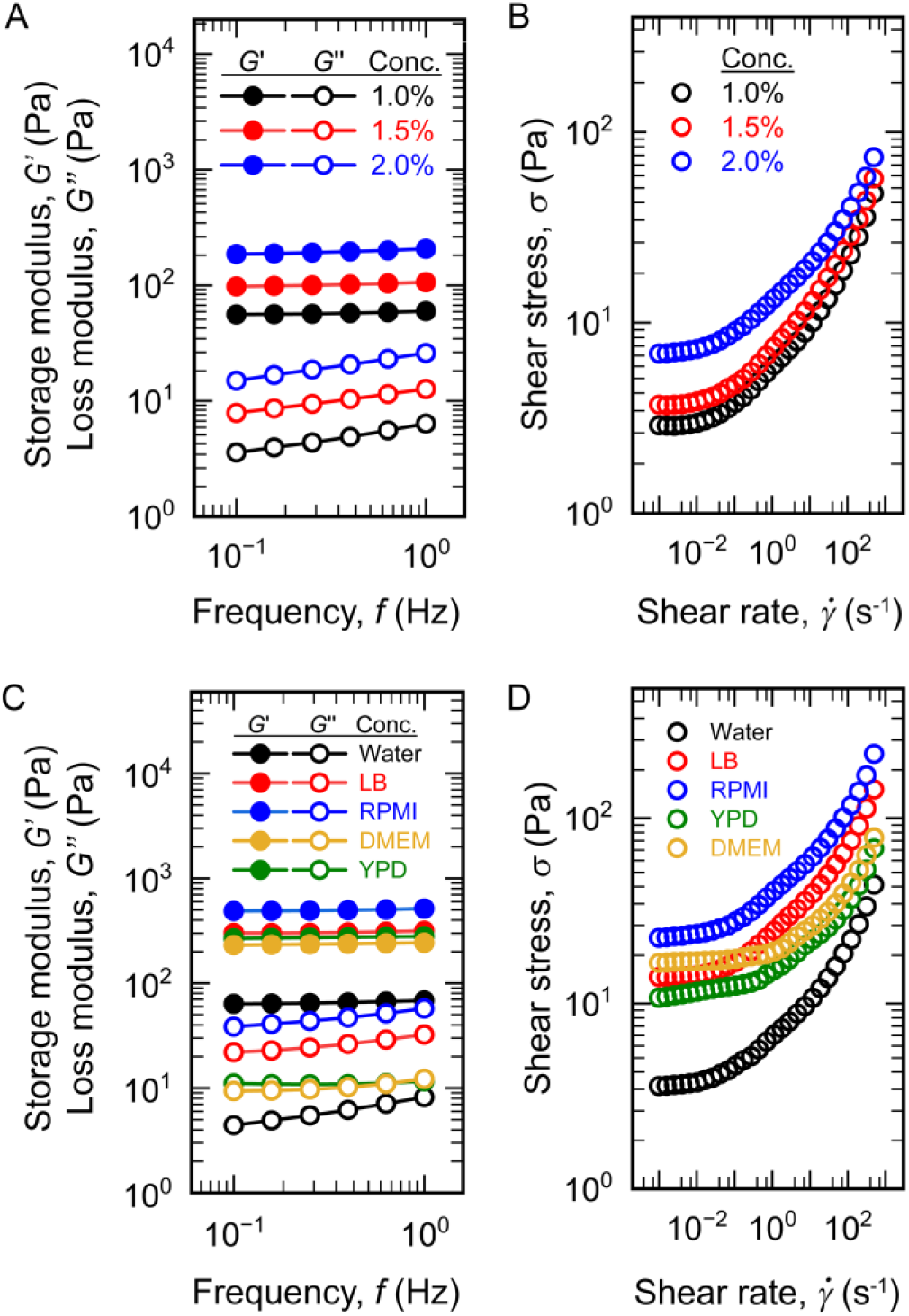
Rheological characterization of jammed agarose microgels. (**A**) A small amplitude oscillatory strain (1%) is applied to the jammed microgel samples over a range of frequencies and shear moduli are recorded. The storage (elastic) moduli, G’, for all samples are both frequency independent as well as larger than the loss (viscous) moduli, G’’. Thus, these jammed microgels behave as soft solids within this range. The average G’ of a jammed microgel system can be altered by the concentration (w/v %) of agarose used. Hence, these microgels exhibit tunable stiffness. (**B**) jammed microgel samples are subjected to unidirectional shear at various rates the shear stress responses are recorded. At high shearing rates, the shear stress is dependent on the shear rate indicating the fluid-like behavior of the microgel packing. At low shear rate, the shear stress is independent of the shear rate which indicates a solid-like behavior. (**C**) and (**D**) We explore similar parameters with microgels prepared in a variety of standard culture media, and observe that the aforementioned properties remain conserved. Importantly, our manufacturing process is not adversely affected by the media composition, hence indicating its broad applicability.

To accommodate easy placement and retrieval of cells as well as cell proliferation, differentiation, and migration in 3D, the growth matrix should have self-healing material properties and allow for easy remodeling of the cellular microenvironment. For instance, a bacterial colony growing under 3D confinement actively deforms the surrounding matrix and expands over time as individual cells continue to proliferate. Similarly, mammalian cells form spheroids in a 3D culture expands in 3D by actively pushing against its surrounding matrix. We test the self-healing material properties of the jammed agarose microgel media using a rheometer; we apply a unidirectional shear at varying shear rates while recording the stress response for each sample. Similar to previously reported jammed microgel systems [56], we find that the packings of agarose microgels are soft solids that can transition between a fluidized state and a jammed solid state as a function of applied shear. Above a threshold shear rate, they transition from soft solid to liquid-like behavior (Fig. 2B, SI Fig. 2C, and SI Fig. 2F).

The ability of jammed agarose microgels to transition back and forth between a solid and fluidized state carries significant implications from an experimental perspective. The yield stress property of jammed agarose microgels permits both spatial and temporal micromanipulations without causing plastic deformations or altering the mechanical properties. This flexibility enables sample seeding, retrieval, region/cell-specific probing, as well as the introduction of additional components such as drugs, signaling cues, cellular markers, etc., during the course of an ongoing experiment itself in a user-defined manner. In rigidly cross-linked polymeric scaffolds, it is not possible to extract cells post-seeding, which precludes possibilities for DNA/RNA/protein extraction or immunostaining without disrupting the bulk structure.

### 3.3 Jammed agarose microgels for mammalian cell culture

To support and grow cells in 3D, the jammed microgel platform requires a cell type-specific growth medium as the continuous phase between the microgel granules. We accommodate this by synthesizing microgels directly inside standard growth media as the continuous phase and evaluating their material properties. Regardless of the chemical nature of the different growth media, the material properties of jammed agarose microgels remain broadly conserved, indicating the universal efficacy of this method (Fig. 2C and 2D). Utilizing the self-healing nature of the jammed microgel media, we directly inject mammalian cells and homogeneously distribute them throughout the 3D media. Furthermore, the jammed microgels prepared with different growth media retain excellent optical transparency (SI Fig. 3A). Leveraging this physical property, we are able to capture high-quality images of living cells expressing a fluorescent reporter (Fig. 3A). Since agarose surfaces do not permit cell adhesion, cells inside the jammed microgels media are held in a 3D suspended state supported by multiple agarose microgel particles (SI Fig. 3B). Additionally, the charge neutrality of agarose eliminates interference with the surface properties and cellular interactions.

**Figure 3:**
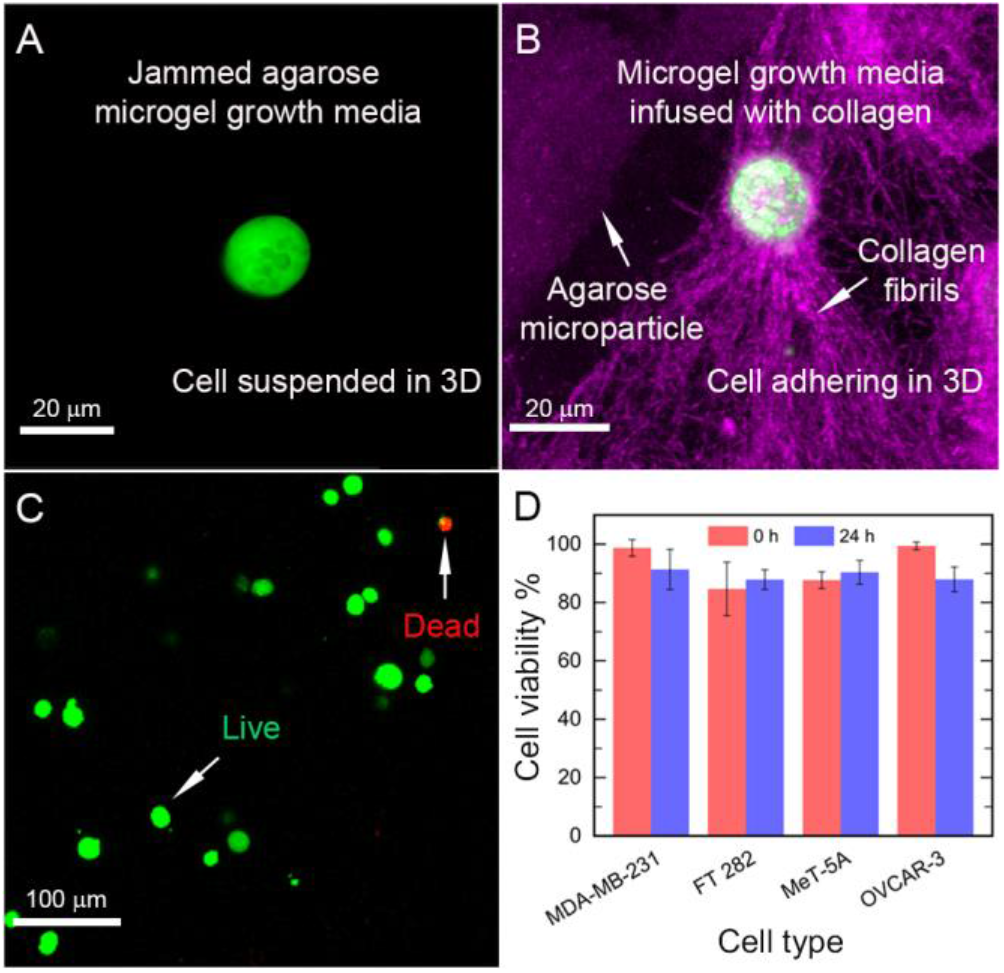
Mammalian cell culture in jammed agarose microgel growth media. (**A**) The optically translucent nature of jammed agarose microgels permit high-quality live imaging of mammalian cells, which remain in a 3D suspended state in absence of any added biopolymer matrices. Pictured here are eGFP-expressing OVCAR-3 cells. (**B**) To promote cellular adhesion, collagen monomers are mixed with the jammed microgel growth medium and are allowed to polymerize within the pores (final concentration of 1 mg/ml). Here an OVCAR-3 cell (green) adhering to and restructuring the polymerized collagen fibrils (purple) inside a jammed agarose microgel medium. (**C**) To evaluate the biocompatibility of these microgels, we homogeneously disperse various types of mammalian cells inside jammed agarose microgels and culture them for 24 hr. We stain for live cells with calcein-AM (green) and dead cells with propidium iodide (red) right after dispersing cells in 3D and 24 hr after culturing in 3D. (**D**) Viability of different cell types measured over 24 hr indicate that jammed agarose microgel systems are biocompatible and can be used as 3D cell growth media.

However, cellular adhesion to biopolymers is critical for both engineering tissue-like mimics and culturing strictly adherent cell types. To culture adherent cells inside a 3D disordered media, we directly polymerize collagen monomers inside the inter-microgel pores of a jammed agarose microgel growth media to achieve a final collagen concentration of 1 mg/ml collagen. The collagen monomers polymerize at 37 °C, creating a network of fibrils within the pore spaces of the microgel. Cells dispersed through the interparticle pores are able to cluster these fibrils by adhering to them and pulling together several of these, indicating local reorganizations of their 3D microenvironment (Fig. 3B).

Although cellular behavior has been extensively studied inside monolithic biopolymer gels of collagen and laminin-rich basement membrane matrix, the disordered nature of jammed microgels with biopolymers inside the inter microgel pore space accurately mimics the tissue microenvironments [57]. Our approach allows the study of individual cells in a confined 3D environment in the presence of extracellular matrices. Furthermore, it provides the capability for region-specific manipulation such as introducing chemical cues or sampling a fraction at different time points by leveraging the self-healing properties of the jammed microgels.

A critical goal for 3D culturing setups is to ensure that cells remain viable over sufficiently long durations. We test the biocompatibility of our jammed agarose microgels by assaying for cellular viability over a 24-hour period (Fig. 3C and 3D). Not only are cells stably maintained within these gels, the non-cancerous pleural mesothelial cell line MeT-5A exhibits robust growth and proliferation over this time course. To rule out any cell type-specific effects, we evaluate a panel of different cells, including epithelial ovarian cancer cells (OVCAR-3), hTERT-immortalized fallopian epithelial cells (FT 282), MeT-5A, and the epithelial breast cancer cells (MDA-MB-231). To meet the requirement of cell type-specific culturing media (RPMI for OVCAR-3, DMEM for the others), we simply change the continuous phase during synthesis to generate microgels compatible with each cell type. These results illustrate the broad applicability of our *in vitro* 3D system for mammalian cell culture.

Promising future directions include culturing organoids, stem cells, and patient-derived primary cell cultures in this matrix. We expect that difficult-to-culture cell types - such as those obtained from *ex vivo* explants - can be maintained with greater success by stiffness matching agarose microgels to their physiological organ/tissue origins. Similar to what has been shown using PEG hydrogels, the capability to make microgels adherent for cells opens up possibilities to generate micro-patterned gel surfaces with essential growth and differentiation factors interspersed at strategic locales[30, 58-63].

### 3.4 Jammed agarose microgels for microbial cell culture

To test the suitability of jammed agarose microgels as a universal biomaterial across different scales, we test the growth and dynamics of yeast and prokaryotes (bacteria) in them. Apart from supporting microorganisms in 3D, jammed microgels also closely mimic their natural habitat. Yeast is widely used to ferment semisolid materials such as dough, whereas bacteria naturally inhabit complex 3D porous environments such as soil, skin pores, gut mucus, and inter-tissue pores. Recently, jammed systems of polyelectrolyte microgels have been utilized for understanding bacterial dynamics within 3D porous media, which has led to a paradigm shift from conventional ideas of motility and spatial interactions[64-67]. However, the coulombic interaction of the polyelectrolyte microgels with the components of the dispersed medium poses a critical challenge and limits the possibility of testing the effect of ionic compounds on microbes in porous media. The jammed agarose microgel media—a charge-neutral granular porous 3D matrix—solves this problem.

To culture yeast in agarose microgel media, we homogeneously disperse a commonly used laboratory strain, *S. cerevisiae*, inside the jammed media. The inter-microgel pore space of the 3D media is filled with Yeast Extract– Peptone–Dextrose (YPD) Medium, which provides a nutrient-rich environment for the yeast cells. Leveraging the optically translucent properties of the agarose microgel medium, we directly visualize the proliferation of yeast colonies using confocal microscopy. This is a consequence of the yield stress nature of jammed microgels – colonies locally and temporarily fluidize the surrounding matrix while growing outward, but they are held in place by the soft solid-like gel (Fig. 4A). To quantify the growth of yeast, we further exploit this optical translucency and use a plate reader to measure the temporal change in absorbance of 600 nm monochromatic light by the overall media. Since, at any time, absorbance is proportional to the total number of cells present, the temporal change in absorbance indicates the growth rate of yeast in the jammed agarose media (Fig. 4B).

**Figure 4:**
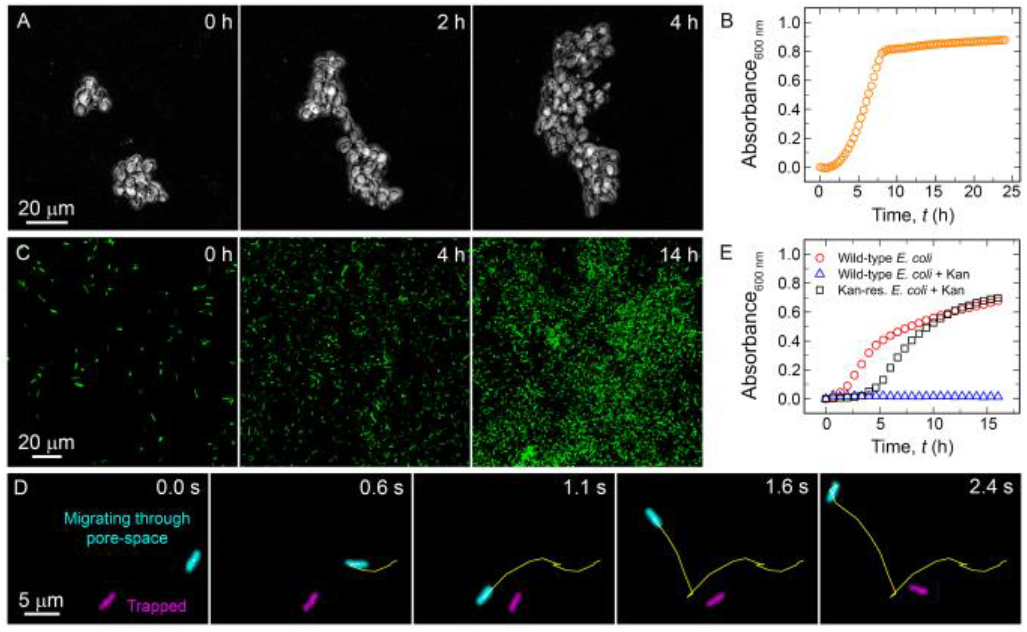
Yeast and prokaryote cell culture in jammed microgel growth media. Our 3D culture media is easily adaptable for a variety of cell types by replacing the continuous phase with a culture media of choice. Here, we show yeast (*S. cerevisiae*) and bacterial (*E. coli*) culture in YPD and LB broth-based jammed agarose microgel. The optically transparent nature of these microgels make them amenable with widely used readout methods such as live-cell imaging – (**A**) for growing *S. cerevisiae* colonies and (**C**) for motile GFP-expressing *E. coli* – as well as (**B**) absorbance-based growth curve assays, as shown for *S. cerevisiae*. It is also possible to maintain a spatially fixed field of view over several hours with these microgels since they behave as soft solids. This is also enabled by the inert and non-charged nature of agarose, which remains stable at standard culturing conditions, thereby permitting long-term live cell imaging and culturing. Our platform is also suitable for studying the dynamics of motile bacteria, such as the wild-type *E. coli* used here. Within the inter-particle pore spaces, individual *E. coli* bacterium exhibit hopping-and-trapping motion (**D**), as previously reported in porous granular media. High-resolution tracking of this behaviour is enabled by the microgel’s optical transparency. The charge neutrality is a critical property of these jammed microgels, which makes them compatible with commonly used charged antibiotics, such as kanamycin. Growth curves in LB-based microgels (**E**) with and without kanamycin using both wild-type and kanamycin-resistant *E. coli* indicate that this polycationic antibiotic retains its antimicrobial activity in jammed agarose microgels.

Similar to yeast cultures, the jammed agarose microgel media—prepared by infusing LB in the inter-microgel pore space—supports the growth of bacterial populations. The growth of *E. coli* populations constitutively expressing GFP is directly imaged using a confocal fluorescence microscope (Fig 4C). However, unlike yeast, *E. coli* cells are motile. While in a homogeneous liquid environment, *E. coli* cells move via run-and-tumble motion, in porous media, the microbial dynamics are governed in large part by the microscale architecture of the system. Recent reports show that these bacteria explore their porous microenvironment by traversing the interconnected pore spaces by hopping—fast-directed motion—between local traps where cellular motion is halted [64, 66, 67]. Similarly, we observe the hopping-and-trapping motility of *E.coli* cells inside the jammed microgel growth media (Fig. 4D) confirming that these cells are proliferating inside a truly disordered 3D media, much similar to their natural habitat.

While the 3D growth media made from jammed polyelectrolyte microgels have been extensively used to study bacterial colony growth [15], the coulombic interactions between the microgels and ionic compounds in the media limit their applicability. One such example is antibiotics-based selection - a commonplace practice in microbiology laboratories due to its relative ease of use and efficacy of action. However, several antibiotic molecules are charged species that can be sequestered by the microgels and hence become ineffective. Since a predominant fraction of microbiological research extensively employs antibiotic selection, this presents a considerable barrier to the broader deployment of jammed polyelectrolyte microgel systems. Indeed, inside a 3D medium prepared by jamming negatively charged microgels (carbomer 980), kanamycin—a polycationic aminoglycoside molecule—when used at a concentration of 50 μg/mL—higher than the minimum inhibitory concentration in liquid LB—fails to halt *E. coli* growth (SI Fig. 4A). While kanamycin is rendered ineffective in charged microgels, it restricts the growth of wild-type *E. coli* in neutral jammed agarose microgels (Fig. 4E). This leads to two significant outcomes – one, that charge-neutral agarose microgels are highly suited for *in vitro* experiments which require external treatments or supplementation with antibiotics, growth factors, drugs, lentiviral vectors, and common post-transfection/transformation selection markers. Secondly, the absence of charge interactions negates the possibility of the active agent being sequestered by the biomaterial itself. This enables a precise, reproducible, and tractable experimental design. Furthermore, this functionality of our charge-neutral microgels has significant implications for investigating selection pressures in a generalized manner in 3D. For instance, substratum stiffness impacts the biofilm formation ability of bacteria [68-73], which may alter drug susceptibility. Phenotypic responses by microbes to variations in surface properties may also be linked to antibiotic resistance mechanisms. From a translational perspective, our experimental setup can help engineer models of complex physiological niches such as tissues. These are often the site of active host-pathogen interactions, wherein selection pressures and adaptive morphologies drastically impact the eventual outcomes.

Agar, agarose, or gelatin have also been extensively employed to encapsulate live bacteria for biomedical applications such as culturing novel microbial strains and efficient *in vivo* delivery of probiotics, as well as to develop high-throughput drug screening platforms[74-79]. In these instances, the microbes are physically constrained within a rigid matrix. We also explore the possibility of encapsulating live GFP-positive *E. coli* cells within the agarose microgel particles. These granules are manufactured using LB broth as both the continuous and dispersed phases. Encapsulated bacteria remain viable and effectively immobilized in 3D space (SI Fig. 4B). Compared to previous attempts, our one-pot synthesis obviates the need for dedicated microfluidic network fabrication, polymeric cross-linking reactions, or extensive washes.

### 3.5 Embedded 3D bioprinting in jammed agarose microgel media

The ability to spatially arrange cells in 3D with precise control over population shape, size, and composition increase the versatility of a 3D cell growth medium. Further, precisely patterning multiple cells in 3D allows for examining intercellular interactions. While a 3D network of biopolymers (e.g., collagen, matrigel, etc.) can support cells in 3D, they do not allow for precise placement of cells in 3D; any insertion of a nozzle through these monolithic matrices creates permanent damage and hence irreversibly alters their mechanical properties. For the same reason, retrieving cells without damaging these monolithic matrices is severely challenging. These challenges are overcome by directly 3D printing cellular constructs inside the jammed agarose microgel systems [26, 28, 34, 37-42, 80-96]. W e exploit the yield stress nature of jammed agarose microgels to precisely 3D print complex cellular structures inside them. In the presence of a shear force, jammed agarose microgels can transition to a fluid-like material. The shear forces generated by an injection nozzle moving through the jammed microgels in a designated pathway locally fluidize the microgel packing and simultaneously inject cells in the fluidized zone. This fluidized zone reverts to a stable, solid-like conformation once the shear is removed, trapping the cells in the desired location in 3D. The print quality and the surface roughness of 3D printed constructs depend on the granularity of the microgels. Jammed agarose microgels with even smaller granules can be produced by utilizing a tissue homogenizer (SI Fig. 5A and SI Fig. 5B). Utilizing this system, we demonstrate the inoculation of bacterial populations of complex geometries inside a 3D growth medium (Fig. 5A).

**Figure 5:**
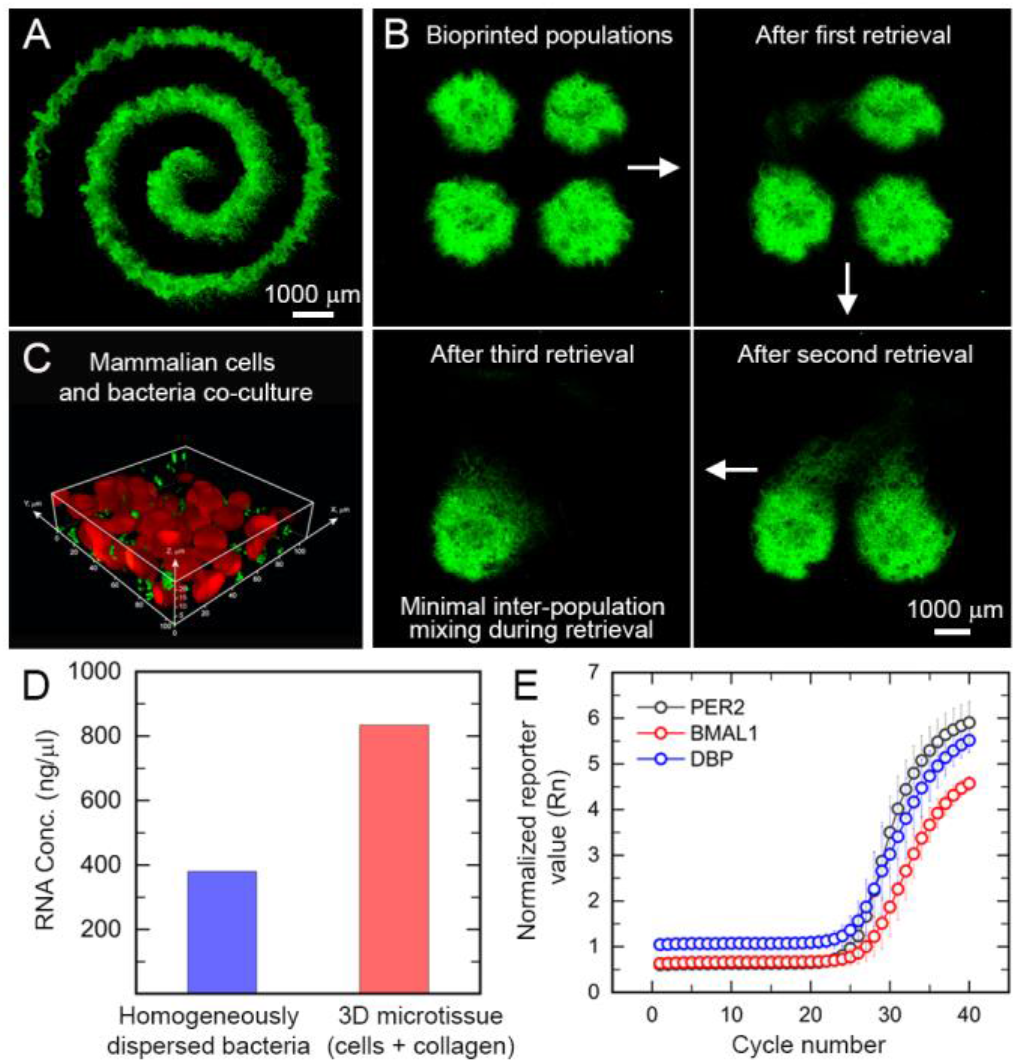
3D bioprinting in jammed agarose microgels. 3D bioprinting is a valuable tool for spatially arranging and manipulating cells as per user-defined conformations. (**A**) As shown using GFP-expressing *E. coli*, the self-healing ability of jammed agarose microgels permits extrusion bioprinting of complex geometries with bioinks. Since these microgels can be locally and reversibly fluidized, individual bacterial populations of GFP-expressing *E. coli* are selectively retrieved (**B**) without perturbing neighbouring structures. Hence, spatiotemporal sampling from an ensemble arrangement is possible without disrupting the bulk gel. This also allows modifications to an existing microgel culture, as shown by (**C**) printing RFP-expressing OVCAR-3 cells into a GFP-expressing *E. coli*-laden microgel bath. The co-culture model allows direct visualization of inter-species interactions using live-cell imaging, since our microgel bath itself is optically transparent. Such co-culture models can be maintained long-term without transferring into a separate culturing setup, since the microgel bath is made in nutrient broth. (**D**) QuBit Broad Range RNA Assay values for samples of RNA extracted from bacterial cultures homogeneously dispersed and grown overnight in a LB-based agarose microgel, as well as microtissues comprising a mixture of cells and collagen injected into a DMEM-based agarose microgel. (**E**) qRT-PCR absorbance curves for cellular genes using RNA extracted from mammalian cells. Our protocols permit direct extraction and quantification of RNA from gel-grown and gel-embedded samples, implying potential applications for investigating differential gene expression in 3D cultures.

The potential for the jammed microgel system is not limited to merely 3D bioprinting by patterning bioinks or introducing multiple cell types. Rather, self-healing material property may be exploited to selectively retrieve parts of a printed assemblage with minimal perturbation to other regions. To test this hypothesis, we 3D print four isolated bacterial populations in close vicinity inside agarose microgel media and sequentially retrieve them (Fig. 5B). Even when the inter-population distance is less than their size, we are able to retrieve each population completely without causing any major perturbations. This is a significant advance over cross-linked 3-D culturing systems - the rigid, monolithic nature of which render such micro-manipulations untenable. Once cell populations can be selectively retrieved from the 3D culture, they can be subjected to standard assays such as metabolomics, proteomics, or whole transcriptome sequencing. To prove this concept, we extract RNA from bacterial colonies cultured in jammed agarose microgel growth media, as well as from microtissues composed of mammalian cells seeded in a collagen matrix and suspended within agarose microgels (Fig. 5D). We also carry out qRT-PCR amplification for three different cellular genes using RNA extracted from 3D microtissue samples (Fig. 5E). Together, these two capabilities significantly expand the boundaries of *in vitro* 3D culturing. The selective retrieval strategy combined with RNA extraction and amplification from captured samples allows one to locally and spatiotemporally sample individual niches within a complex system without disrupting the global architecture. By incorporating micro-scale mechanical manipulators [97], this can be narrowed down to the single-cell level.

A universal 3D cell growth medium with the option of 3D bioprinting cellular constructs paves the pathway to a systematic exploration of intercellular interactions. Specifically, in the context of infection biology, it has been a longstanding goal to develop physiologically relevant *in vitro* setups to study disease progression. While animal models are vastly superior, they pose challenges to live cell imaging, tissue-level observations in real-time, and standardized control over the experimental dynamics. Co-culture systems using transwells, organoids, and microfluidic setups have tried to address this by incorporating multiple cell types or providing 3D-like culturing conditions, which cannot be realized using conventional monolayer cultures [98]. However, these scaffolds either constrain the interacting populations to an adherent/liquid-suspended state or utilize constrained geometries and flow-based networks, which may disrupt the structural integrity and limit the scale of tissue-like multicellular aggregates. The 3D nature and self-healing ability of agarose microgels help overcome these shortcomings. Briefly, they support cells in a suspended 3D state while allowing temporally separated seeding of distinct cell types within the same system. Additionally, the bath dimensions can be readily adjusted based on the desired culturing scale. Here, we demonstrate a co-culture model of OVCAR-3 cells printed within an *E. coli*-laden RPMI-based jammed agarose microgel bath. Our optically transparent microgel allows real-time imaging of such a system (Fig. 5C and Supplementary Video). These *in vitro* setups would help model processes such as bacterial or cancer cell invasion of epithelial layers and immune cell dynamics in response to pathogens.

## 4. Conclusion

Existing microgel-based 3D culturing platforms have vastly expanded our ability to interrogate cellular organization and interactions in settings mimicking physiological niches. These systems have been used to arrange, maintain, probe, and manipulate cells in well-defined microenvironments [97]. Combining these hydrogels with 3D bioprinting, an impressive range of structural features has also been rendered. However, in-house fabrication of these microgels involves multi-step synthesis protocols [39, 80, 99], which substantially increases both preparation time and precautions required to maintain sterility. In contrast, many commercially available microgels [29, 56] are made from negatively charged polyelectrolyte hydrogels that interfere with the bivalent cations of the cell growth media [32]. Additionally, maintaining the sterility of commercially available microgels remains a persistent problem; several microgels are not compatible with either UV or heat sterilization, two of the most widely adopted practices in cell culture laboratories. A 3D medium made of packings of agar/agarose microgels solves this problem. Prior efforts at making agar microgels have used phase separation techniques by introducing either oil or PEG as an emulsifying agent to break up an aqueous solution of molten agarose into discrete particles [100-103]. However, these either involve dedicated microfluidic networks or require extensive wash steps which do not entirely remove traces of the foreign agent. Our approach overcomes these limitations by employing a simple bench-top setup, commonly available mixing equipment, and a one-step synthesis protocol without any additional chemical intermediates or washing steps, which drastically reduces potential chances for contamination. The agarose itself is heat sterilized prior to microgel manufacturing with the additional flexibility for UV sterilization post-manufacture. The ease and simplicity of synthesis put it within reach of most experimental laboratories without any requirement for specialized training or sophisticated equipment.

Jammed agarose microgels provide a truly universal 3D platform that can simultaneously host diverse cell types and inter-species co-cultures, enable high-quality live cell imaging, be functionally compatible with commonly used selection markers, allow 3D bioprinting with live bioinks, and permit localized sampling. Our study hence lays the groundwork for a highly accessible and versatile 3D culturing platform. Both the modular synthesis process and the inert nature of agarose contribute to the versatility. The non-reactive property is important for live-cell encapsulation without affecting the surface properties of the cells, since agarose does not carry residual charges. The charge neutrality of this polymer also allows antibiotics-based selection, unlike many existing 3D platforms which use charged microgel particles. The distinct advantages in terms of chemical inertness and charge neutrality greatly expand its applicability for high-throughput screening applications. It is also possible to incorporate selective experimental cues - for instance, pharmacological agents and growth/differentiation factors – while following existing protocols designed for conventional 2D systems.

Jammed microgels also serve as scaffolds to assemble complex, multicellular, tissue-mimicking structures. This is made possible by extrusion bioprinting, which confers the ability to spatially arrange cells in 3D. Designing tissue-like mimics is feasible using our scaffold since the bulk gel can be modified to make it suitable for adherent cell culture. In a significant advance over several existing rigid 3D scaffolds, we can spatiotemporally probe and retrieve samples without perturbing or compromising the structural integrity of the system. This feature, in combination with our capability to extract RNA from agarose microgel samples, suggests possibilities to study differential gene expression in 3D cell culture models. With further optimizations, this could likely be extended to cover the transcriptome, proteome, and metabolome domains as well. Hence, we envision extremely promising avenues for further advances in 3D cell culture, bioprinting, and tissue bioengineering using jammed agarose microgels.

The disordered, porous, and granular architecture of jammed agarose microgels mimics scenarios frequently encountered by soil and gut mucus-resident bacteria, which navigate porous environments that are structurally similar to polymeric microgels. This charge-neutral matrix also opens up future opportunities to study electrotaxis in 3D. Furthermore, since the shear moduli of individual agarose granules is of the order of ∼10 kPa, they can withstand shear forces due to fluid flow. Thus, the solid packings of agarose granules can be easily housed in confined chambers, and leveraging the transparent nature of fluid flow through the inter-microgel pores can be easily visualized. Hence, the jammed agarose microgels can be used to study bacterial migration through a porous medium under flow and the transition of planktonic cells to form biofilms under local confinement. Similarly, by flowing nutrient-rich medium bioprinted cellular constructs can be maintained for long-term inside jammed microgel growth media. Finally, we envision that by flowing selective morphogens through the jammed agarose microgels, 3D bioprinted constructs can be differentiated to create multifunctional organoids.

## Author contributions

TB designed and conceptualized the overall project. RB provided essential guidance and support for cell viability studies. SM, with critical help from MG, performed all the experiments. All authors participated in writing the manuscript.

## Acknowledgments

We acknowledge Prof. Robert Austin for providing us with the *E. coli* cells, Dr. Shashi Thutupalli for giving us the yeast cells, and Dr. Shaon Chakrabarti for providing 3T3 cells, as well as qRT-PCR protocols and reagents. We are grateful to Dr. H. Krishnamurthy and the Central Imaging and Flow Cytometry Facility at NCBS. We also acknowledge the NCBS common equipment facility for arranging the Anton Parr MCR 302e rheometer and NextGen sequencing facility for arranging Qubit. We thank Ashitha B Arun, Jimpi Langthasa, Shyamili Goutham, and Mallar Banerjee for sharing cell culture protocols as well as for assistance with the experiments. TB acknowledges intramural research grant from NCBS. SM acknowledges personal support through NCBS GS program. RB acknowledges support from the Wellcome Trust/DBT India Alliance (IA/I/17/2/503312) and the Department of Biotechnology, India (DBT) (BT/909 PR26526/GET/119/92/2017).

## Supplementary Information

**SI Figure 1:**
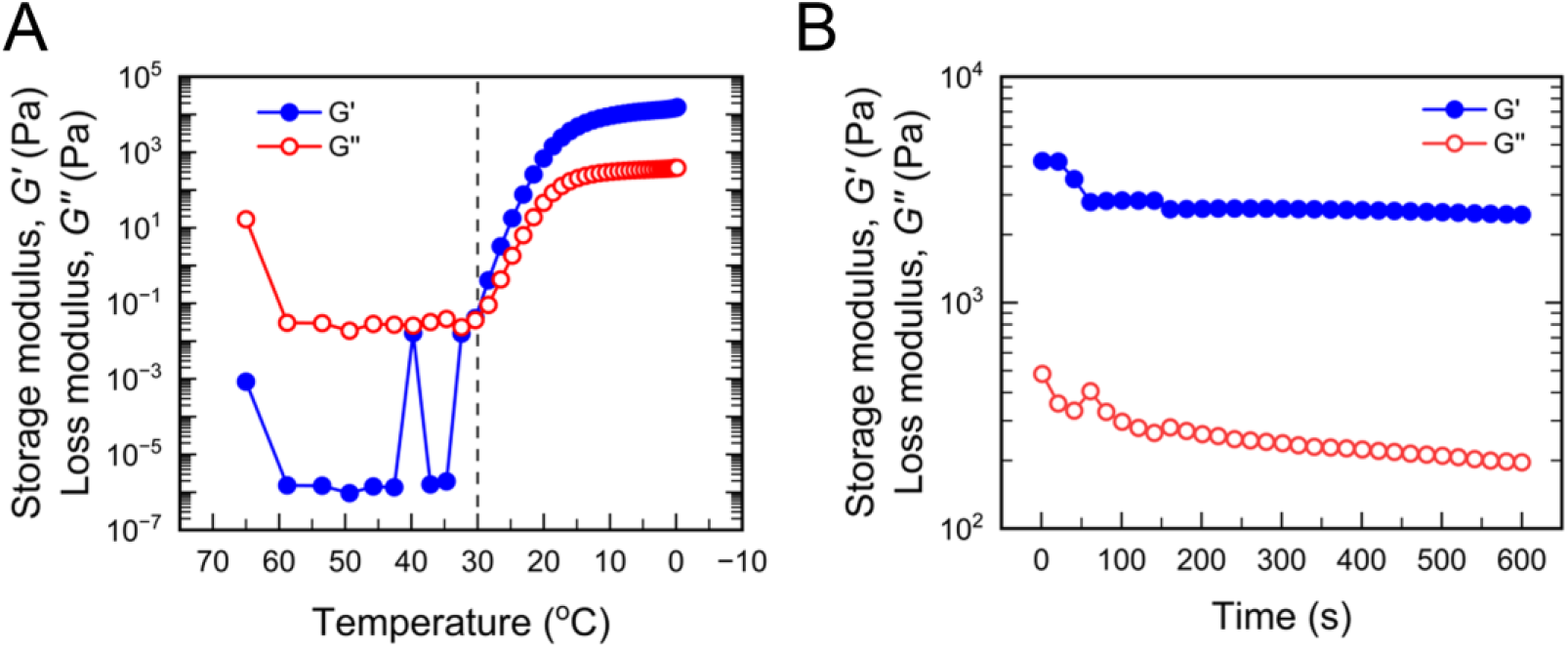
Gelation dynamics of molten agarose. (**A**) We characterize the gelation temperature for a molten solution of 1% agarose in water, wherein G’’ > G’ signifies the liquid (molten) state, and the crossover point where G’ > G’’ indicates gelation of the agarose into a solid. We find that 1% molten agarose commences solidification at 30°C, far higher than the typical temperatures maintained in the ice bath-cooled continuous phase employed in our manufacturing process. (**B**) We study the rate of gelation at 0°C for molten 1% agarose maintained initially at 65°C. The solidification is instantaneous at this temperature difference, as indicated by the persistent G’ > G’’ trend.

**SI Figure 2:**
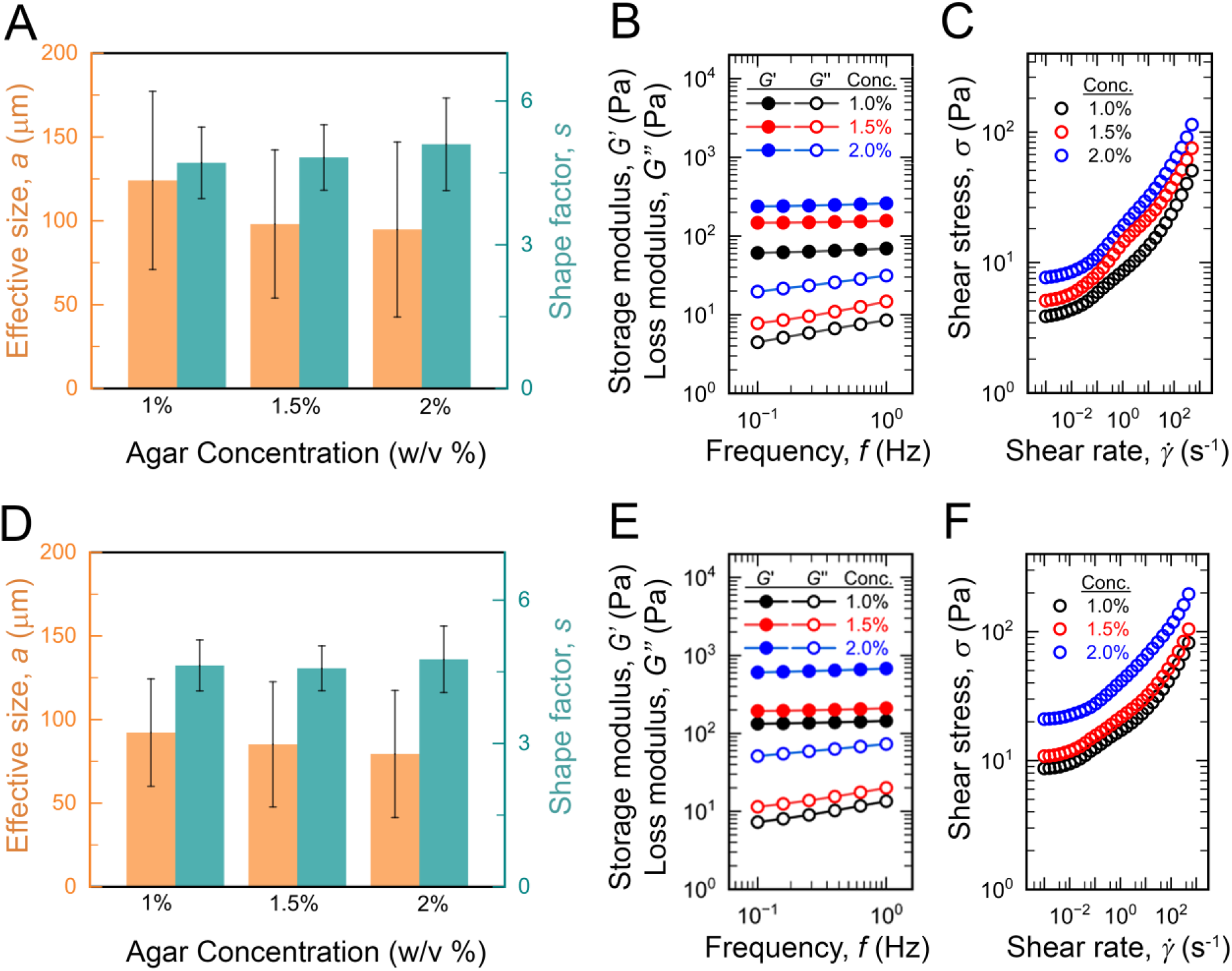
Characterization of agar microgels. Using the manufacturing process described for agarose microgels, we also synthesize and characterize jammed microgels prepared using (**A, B**, and **C**) Sigma and (**D, E**, and **F**) Qualigens agar. (**A** and **D**) We observe a consistent particle size distribution, which is also quite similar to that obtained using agarose. (**B** and **E**) There is no time-dependent rheological behaviour for these microgels, which behave as soft solids at low shear stress regimes, as evidenced by small (1%) amplitude oscillatory shear over different frequencies. (**C** and **F**) Both these agar-based microgels also exhibit yield-stress behaviour, transitioning from a liquid-like to solid-like nature under low shear rates.

**SI Figure 3:**
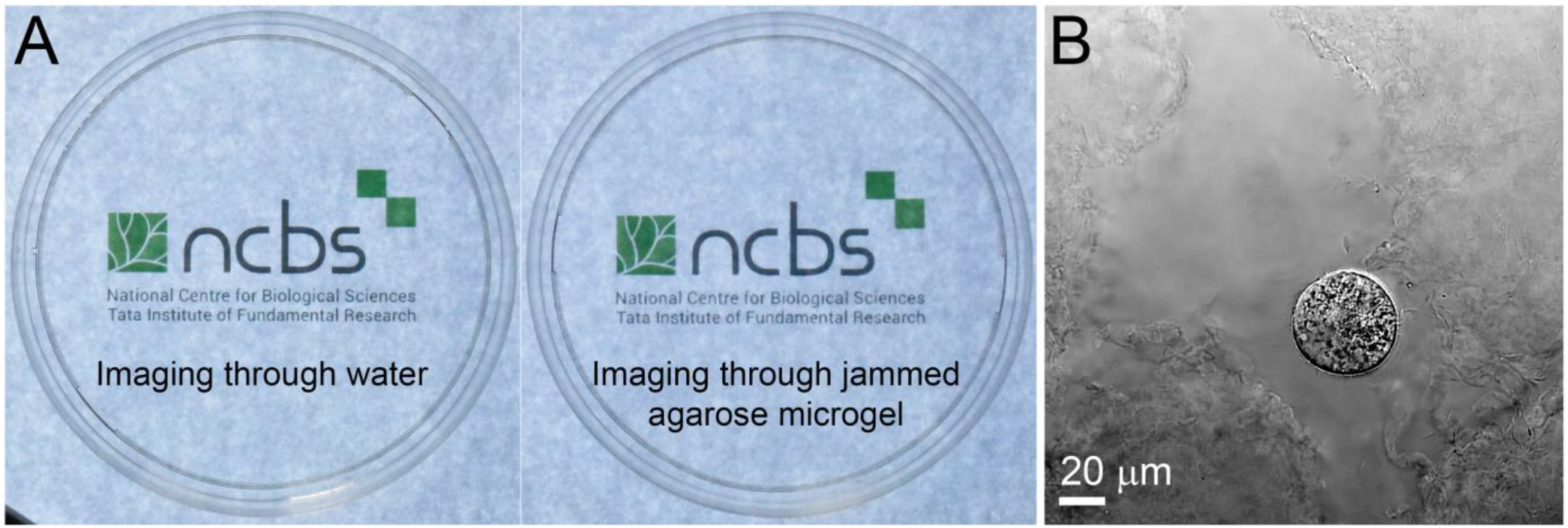
Optical transparency of jammed agarose microgels for imaging applications. (**A**) The NCBS institutional logo viewed through (*left*) water and (*right*) agarose microgels prepared using water as a continuous phase, demonstrating the latter’s high degree of transparency. (**B**) Brightfield image of a single OVCAR-3 cell suspended between several agarose microparticles within a jammed agarose microgel. Not only are cells stably cradled within a suspended environment, we can clearly visualize their 3D morphology within this scaffold using both brightfield and fluorescence microscopy.

**SI Figure 4:**
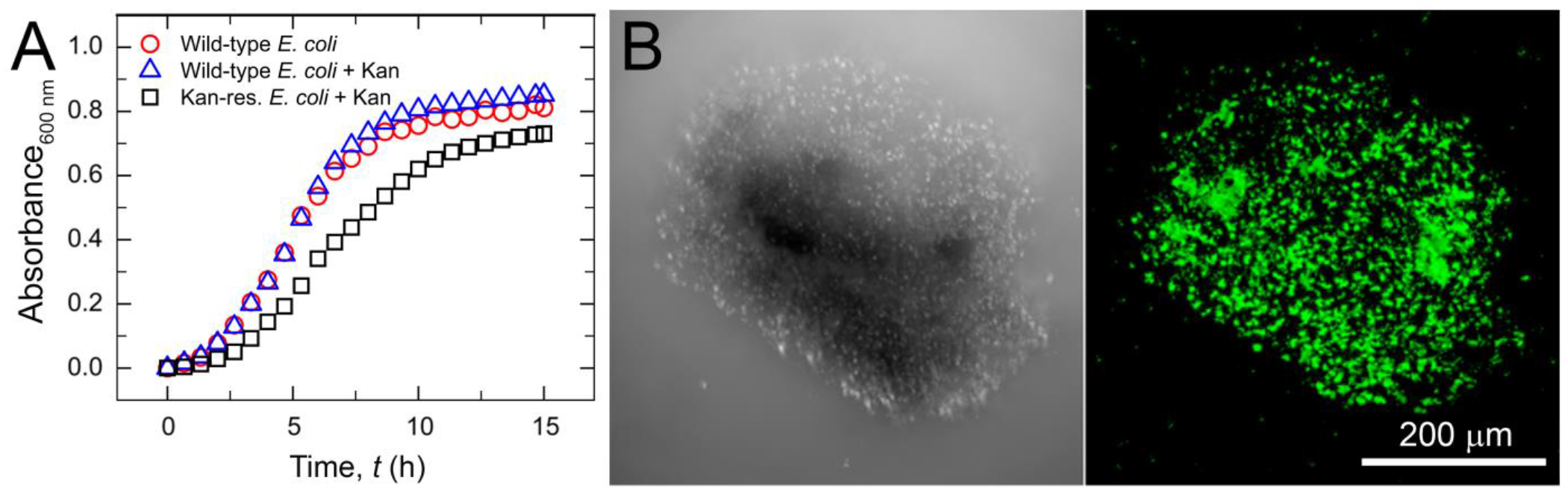
Multi-functionality of jammed agarose microgels. (**A**) 3D growth media manufactured using negatively charged polymeric particles sequester polycationic antibiotics such as kanamycin, rendering them ineffective against bacterial growth. As seen here, both wild-type and kanamycin-resistant *E. coli* strains achieve substantial growth in the presence or absence of kanamycin within these charged microgels. Hence, such platforms cannot be used to study the effects of charged small molecules. (**B**) Live GFP-expressing *E. coli* under monolithic confinement within agarose, manufactured using the same flash solidification method as described for generating agarose microgels. The modularity of the synthesis process enables such diverse capabilities without any significant alteration to the workflow.

**SI Figure 5:**
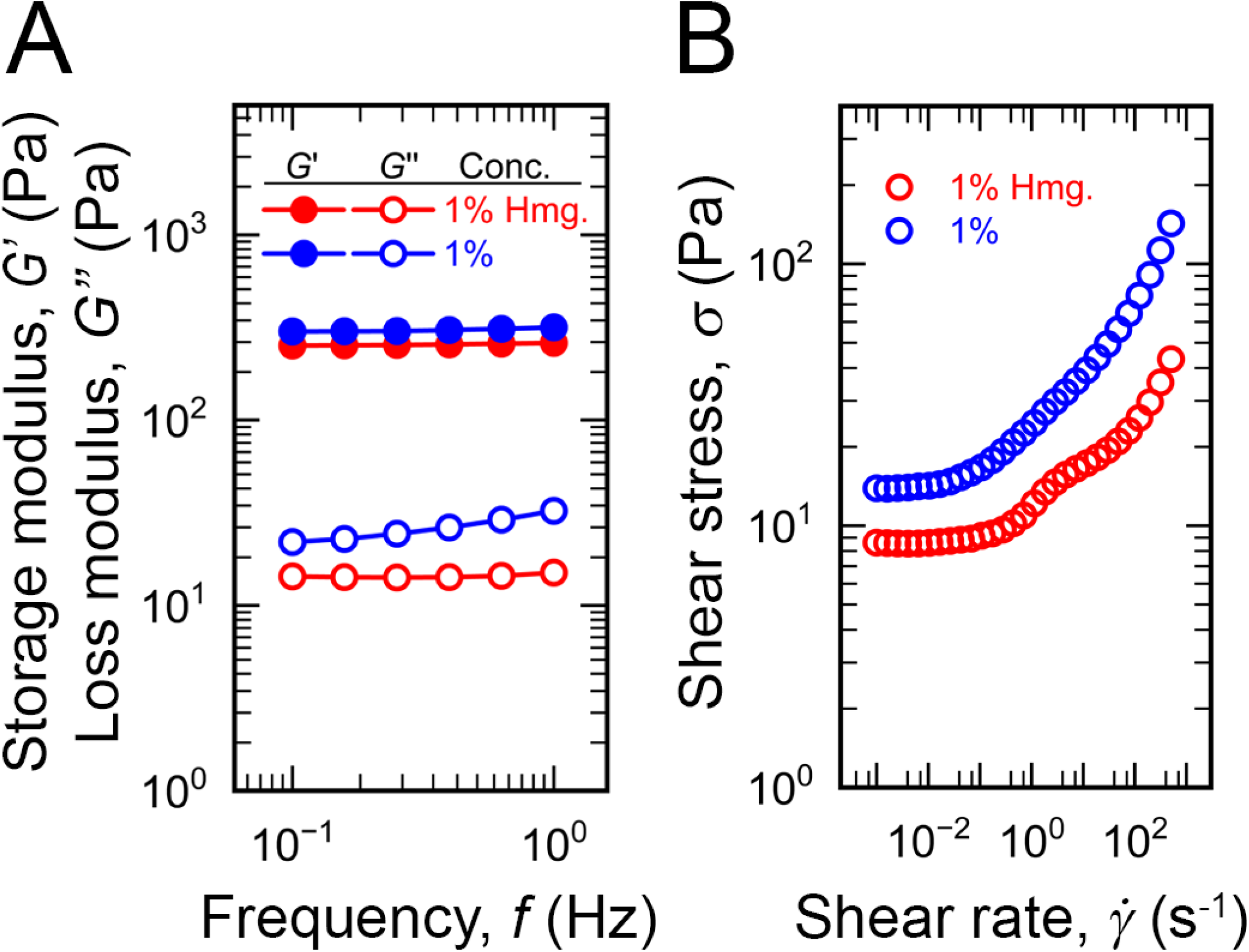
Embedded 3D bioprinting applications in jammed agarose microgel baths. (**A**) Low (1%) amplitude frequency sweeps and (**B**) resultant shear stress responses for varying rates of unidirectional shear on agarose microgels manufactured with LB as the continuous phase, with and without homogenization. The smaller particle sizes resulting from homogenization make for a softer, lower yield-stress matrix, which enables 3D bioprinting with finer resolution.

